# *Trochodendron aralioides*, the first chromosome-level draft genome in Trochodendrales and a valuable resource for basal eudicot research

**DOI:** 10.1101/650424

**Authors:** Joeri S. Strijk, Damien D. Hinsinger, Feng-Ping Zhang, KunFang Cao

**Affiliations:** State Key Laboratory for Conservation and Utilization of Subtropical Agro-bioresources, College of Forestry, Guangxi University, Nanning, Guangxi 530005, China; Biodiversity Genomics Team, Plant Ecophysiology & Evolution Group, Guangxi Key Laboratory of Forest Ecology and Conservation, College of Forestry, Daxuedonglu 100, Nanning, Guangxi, 530005, China; Alliance for Conservation Tree Genomics, Pha Tad Ke Botanical Garden, PO Box 959, 06000 Luang Prabang, Laos; Evolutionary Ecology of Plant Reproductive Systems Group, Kunming Institute of Botany, Kunming, China

**Keywords:** Trochodendron aralioides, chromosome-level genome assembly, Hi-C assembly, basal eudicot

## Abstract

**Background:** The wheel tree (*Trochodendron aralioides*) is one of only two species in the basal eudicot order Trochodendrales. Together with *Tetracentron sinense*, the family is unique in having secondary xylem without vessel elements, long considered to be a primitive character also found in *Amborella* and Winteraceae. Recent studies however have shown that Trochodendraceae belong to basal eudicots and demonstrate this represents an evolutionary reversal for the group. *Trochodendron aralioides* is widespread in cultivation and popular for use in gardens and parks.

**Findings:** We assembled the *T. aralioides* genome using a total of 679.56 Gb of clean reads that were generated using both PacBio and Illumina short-reads in combination with 10XGenomics and Hi-C data. Nineteen scaffolds corresponding to 19 chromosomes were assembled to a final size of 1.614 Gb with a scaffold N50 of 73.37 Mb in addition to 1,534 contigs. Repeat sequences accounted for 64.226% of the genome, and 35,328 protein-coding genes with an average of 5.09 exons per gene were annotated using *de novo*, RNA-seq, and homology-based approaches. According to a phylogenetic analysis of protein-coding genes, *T. aralioides* diverged in a basal position relatively to core eudicots, approximately 121.8-125.8 million years ago.

**Conclusions:** *Trochodendron aralioides* is the first chromosome-scale genome assembled in the order Trochodendrales. It represents the largest genome assembled to date in the basal eudicot grade, as well as the closest order relative to the core-eudicots, as the position of Buxales remains unresolved. This genome will support further studies of wood morphology and floral evolution, and will be an essential resource for understanding rapid changes that took place at the base of the Eudicot tree. Finally, it can serve as a valuable source to aid both the acceleration of genome-assisted improvement for cultivation and conservation efforts of the wheel tree.

## Data description

### Introduction of T. aralioides

The Trochodendraceae family (order Trochodendrales) includes only two species (*Trochodendron aralioides* and *Tetracentron sinense*), both of whom are commercially used and widely cultivated. *Trochodendron aralioides* (or wheel tree) is a native species of the forests of Japan (Honshu – southwards from Yamagata Prefecture, Shikoku, Kyushu, Ryukyu Islands) and Taiwan. Although its hardiness extends to lower temperatures, it is generally restricted to lower temperate montane mixed forests between 600-1700m in Japan. In Taiwan, the range is more extensive, occurring in broad-leaved evergreen forest (2000-3000m) in the central mountain ranges and in northern Taiwan between 500-1250m forming monotypic stands [1]. Over the past century, it has been repeatedly reported from Korea [2–10] although these occurrences are not confirmed in online repositories (e.g. http://plantsoftheworldonline.org/). Properties of *Trochodendron* (e.g. mild-warm temperate range, restricted elevational intervals and natural occurrence in small discontinuous populations) make it difficult to predict the sensitivity of *T. aralioides* to the effects of projected changes in climate [11]. The fossil record shows both species diversity and distribution of the family were much more extensive and continuous during the Eocene (50-52 Ma) to Miocene [12,13]. Unique for basal eudicots, Trochodendraceae have secondary xylem without vessel elements, a property only found in *Amborella* and Winteraceae [14]. This raises interesting questions on the biological conditions or triggers giving rise to such anatomical reversals, and the evolutionary and ecological consequences inherent in them.

Here, we constructed a high-quality chromosome-level reference genome assembly for *T. aralioides* using long reads from the PacBio DNA sequencing platform and a genome assembly strategy taking advantage of the Canu assembler [15]. This assembly of *T. aralioides* genome is the first chromosome-level reference genome constructed for the Trochodendrales order, and the closest relative to core eudicots sequenced to date. The completeness and continuity of the genome will provide high quality genomic resources for studies on floral evolution and the rapid divergence of eudicots.

### Genomic DNA extraction, Illumina sequencing and genome size estimation

High-quality genomic DNA was extracted from freshly frozen leaf tissue of *T. aralioides* (Figure 1) using the Plant Genomic DNA Kit (Tiangen Biotech Co., Ltd), following manufacturer instructions. After purification, a short-insert library (300∼350 bp) was constructed and sequenced on the Illumina NovaSeq platform (Illumina Inc., San Diego, CA), according to manufacturer outlines. A total of ∼124.6 Gb of raw reads were generated.

**Figure 1.**
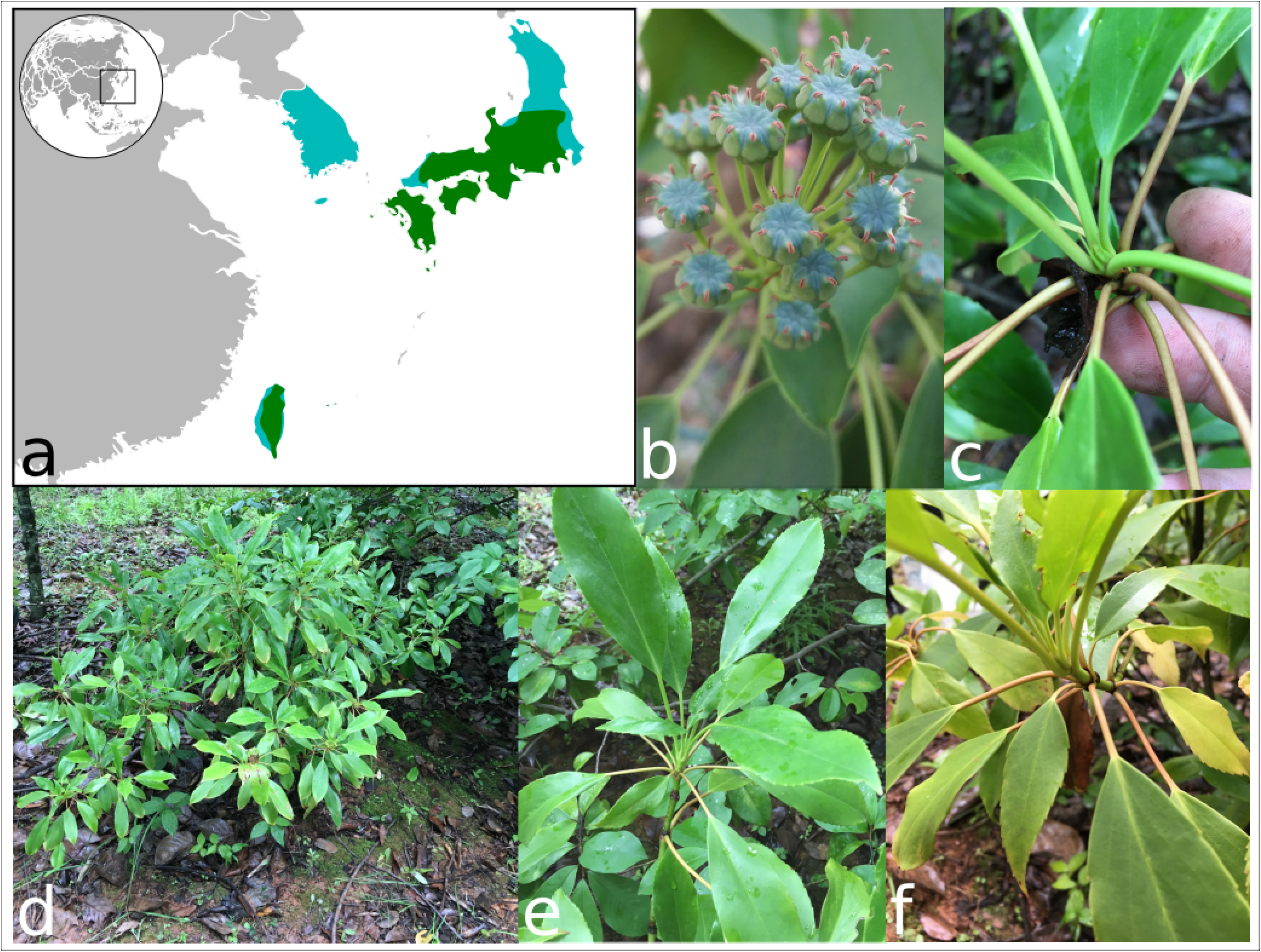
*Trochodendron aralioides* description. a) Geographic distribution. light blue: occurrence according to the Flora of China (at country level), green: occurrence according to GBIF; b) flowers; c) bud; d) general habit; e-f) stem and sprouting bud showing the wheel-like organisation of leaves.

Sequencing adapters were then removed from the raw reads and reads from non-nuclear origins (chloroplast, mitochondrial, bacterial and viral sequences, etc.), screened by aligning them to the nr database (NCBI, http://www.ncbi.nlm.nih.gov) using megablast v2.2.26 with the parameters ‘ -v 1 - b 1 -e 1e-5 -m 8 -a 13’; The in-house script *duplication_rm.v2* was used to remove the duplicated read pairs; low-quality reads were filtered as follows:

1. reads with ≥10% unidentified nucleotides (N) were removed;
2. reads with adapters were removed;
3. reads with >20% bases having Phred quality <5 were removed;

After the removal of low-quality and duplicated reads, ∼124.3 Gb of clean data (Supplementary Table S1) were used for the genome estimation.

The k-mer peak occurred at a depth of 51 (Figure 2), and we calculated the genome size of *T. aralioides* to be 1.758 Gb with an estimated heterozygosity of 0.86%, with a repeat content of 69.31%. This estimate is slightly smaller than previously reported based on cytometry estimate (1.868 Gb) [16]. The GC content was 39.58% (Supplementary Figure S1). A first genome assembly was approximately 1.324 Gb total length, with a contig N50 of 740 bp and a scaffold N50 of 1.079 kb using the Illumina data and the assembly program SOAPdenovo [17]. This first attempt to assemble the wheel tree genome was of low-quality, likely due to its high genomic repeat content and high heterozygosity level.

**Figure 2.**
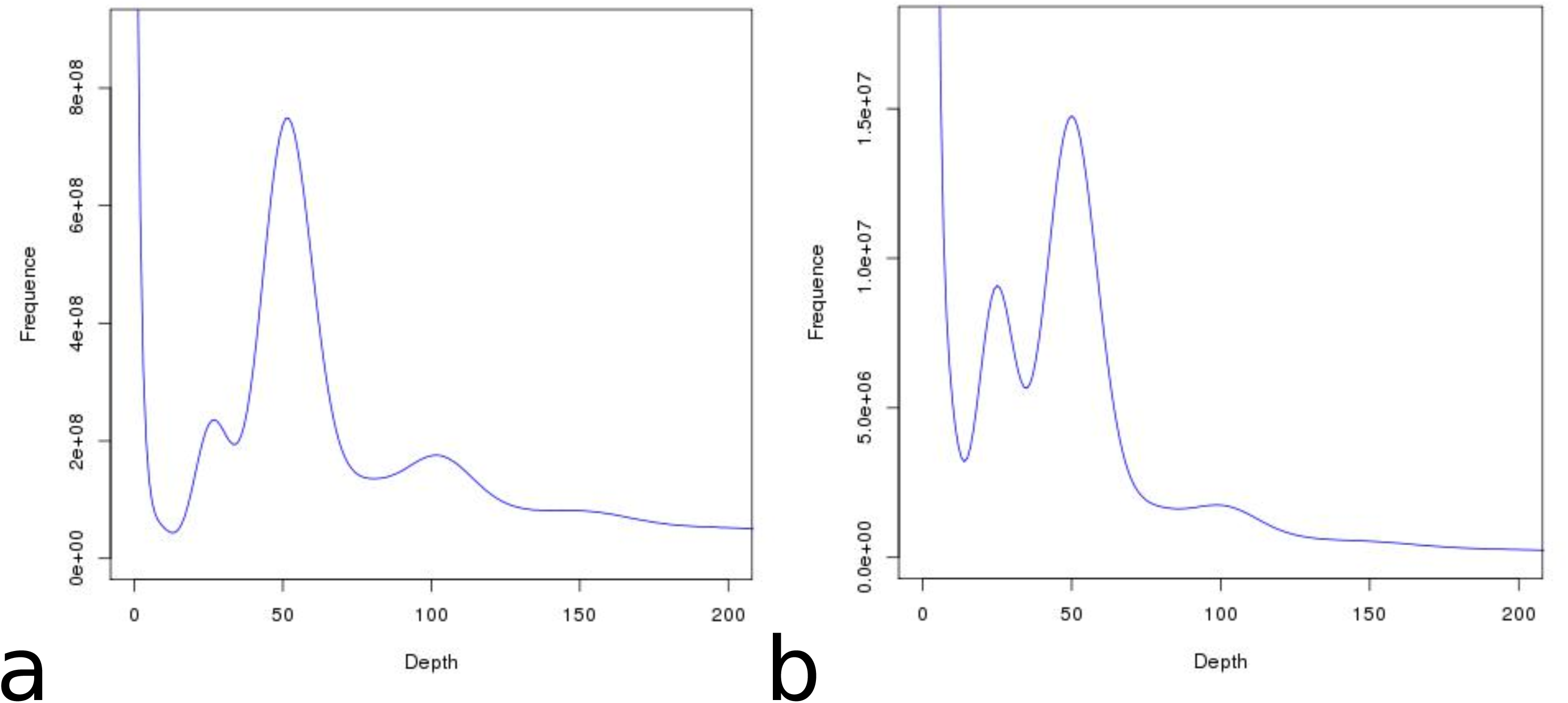
*k*-mer distribution of the *T. aralioides* genome. a) K-mer depth and number frequency distribution; b) K-mer depth and K-mer species number frequency distribution.

### PacBio sequencing

High molecular weight Genomic DNA was sheared using a g-TUBE device (Covaris, Brighton, UK) with 20kb settings. Sheared DNA was purified and concentrated with AmpureXP beads (Agencourt, Bioscience Corp., Beverly, MA) and then used for Single-Molecule Real Time (SMRT) bell sequencing library preparation according to manufacturer’s protocol (Pacific Biosciences; 20-kb template preparation using BluePippin size selection). Size selected and isolated SMRT bell fractions were purified using AmpureXP beads (Agencourt, Bioscience Corp., Beverly, MA) and these purified SMRT bells were finally used for primer-and polymerase (P6) binding according to manufacturer’s protocol (Pacific Biosciences). DNA-Polymerase complexes were used for MagBead binding and loaded at 0.1nM on-plate concentration in 35 SMRT cells. Single-molecule sequencing was performed on a PacBio Sequel platform, yielding a total of 177.80 Gb filtered polymerase read bases (Supplementary Table S1).

### 10X Genomics sequencing

Libraries were built using a Chromium automated microfluidic system (10X Genomics, San Francisco, CA) that allows the combination of the functionalized gel beads and high molecular weight DNA (HMW gDNA) with oil to form a ‘Gel bead in emulsion (GEM)’. Each GEM contains ∼10 molecules of HMW gDNA and primers with unique barcodes and P5 sequencing adapters. After PCR amplification, P7 sequencing adapters are added for Illumina sequencing. Data were processed as follow: Firstly, 16 bp barcode sequences and the 7bp random sequences are trimmed from the reads, as well as low quality pairs. We generated a total of 186.95 Gb raw data, and 183.52 Gb clean reads (Supplementary Table S1).

### Hi-C sequencing data

To build a Hi-C library [18], nuclear HMW gDNA from *Trochodendron aralioides* leaves was cross-linked, then cut with the DPNII GATC restriction enzyme, leaving pairs of distally located but physically interacted DNA molecules attached to one another. The sticky ends of these digested fragments were biotinylated and then ligated to each other to form chimeric circles. Biotinylated circles, that are chimeras of the physically associated DNA molecules from the original cross-linking, were enriched, sheared and sequenced on an Illumina platform as described above. After adapter removal and filter of low quality reads, a total of 193.90 Gb clean Hi-C reads. Sequencing quality assessment is shown in Supplementary Table S2.

### *De novo* Genome assembly

Short PacBio reads (<5kb) were first used to correct the PacBio long-reads using the ‘daligner’ option in FALCON [19], and to generate a consensus sequence. Following this error correction step, reads overlap was used to construct a directed string graph following Myers’ algorithm. Contigs were then constructed by finding the paths from the string graph. Error correction of the preceding assembly was performed using the consensus–calling algorithm Quiver (PacBio). Reads were assembled and error-corrected with FALCON and Quiver to generate 4226 contigs with a contig N50 length of 702 Kb and total length of 1.607 Gb.

FragScaff [20] was used for 10X Genomics scaffolding, as follows:

1. Linked reads generated using the 10X Genomics library were aligned with BOWTIE v2 [21] against the consensus sequence of the PacBio assembly, to obtain Super-Scaffolds; 2) With increasing distance to consensus sequence, the number of linked reads supporting scaffolds connection will decrease. Consensus sequences without linked read supports were then filtered and only the consensus sequence supported by linked reads was used for the subsequent assembly. FragScaff scaffolding resulted in 1469 scaffolds, with a scaffold N50 length of 3.38 Mb.

To assess the completeness of the assembled *Trochodendron aralioides* genome, we performed a BUSCO analysis by searching against the plant universal bench marking single-copy orthologs (BUSCOs, version 3.0) [22]. Overall, 91.4% and 2.8% of the 1440 expected genes were identified in the assembled genome as complete and partial, respectively (Supplementary Table S3). Overall, 94.2% (1,356) genes were found in our assembly. We also assessed the completeness of conserved genes in the *T. aralioides* genome by CEGMA (Core Eukaryotic Genes Mapping Approach [23]). According to CEGMA, 232 conserved genes in *T. aralioides* were identified which have 93.55% completeness compared to the sets of CEGMA (Supplementary Table S3).

The Hi-C clean data were aligned against the PacBio reads assembly using BWA [24]. Only the read pairs with both reads aligned to contigs were considered for scaffolding. For each read pair, its physical coverage was defined as the total bp number spanned by the sequence of reads and the gap between the two reads when mapping to contigs. Per-base physical coverage for each base in the contig was calculated as the number of read pairs’ physical coverage it contributes too. Misassembly can be detected by the sudden drop in per-base physical coverage in a contig.

Following the physical coverage of the resulting alignment, any misassemblies were split to apply corrections. Using the clustering output, the order and orientation of each contig interaction was assessed on intensity of contig-interaction and the position of the interacting reads. Combining the linkage information and restriction enzyme site, the string graph formulation was used to construct the scaffold graph using LACHESIS [25], and the 1469 scaffolds of our draft genome were clustered to 19 Chromosomes (Supplementary Table S4). The *Trochodendron aralioides* genome information is summarized in Supplementary Tables S5.

### Repeat sequences in the wheel-tree genome

Transposable elements in the genome assembly were identified both at the DNA and protein level. RepeatModeler [26], RepeatScout [27] and LTR_FINDER [28] were used to build a *de novo* transposable element library with default parameters. RepeatMasker [29] was used to map the repeats from the *de novo* library against Repbase [30]. Uclust [31] was then used with to the 80-80-80 rule [32] to combined results from above software. At the protein level, RepeatProteinMask in the RepeatMasker package was used to identify TE-related proteins with WU-BLASTX searches against the transposable element protein database. Overlapping transposable elements belonging to the same type of repeats were merged.

Repeat sequences accounted for 64.2% of the *T. aralioides* genome, with 57.2% of the genome identified from the *de novo* repeat library (Table 2). Approximately 53.2% of the *T. aralioides* genome was identified as LTR (most often TEs). Among them, LTRs were the most abundant type of repeat sequences, representing 53.249% of the whole genome. Long interspersed nuclear elements (LINEs) and DNA transposable elements repeats accounted for 0.837% and 2.416% of the whole genome, respectively (Table 2, Supplementary Figure S2).

**Table 2.**
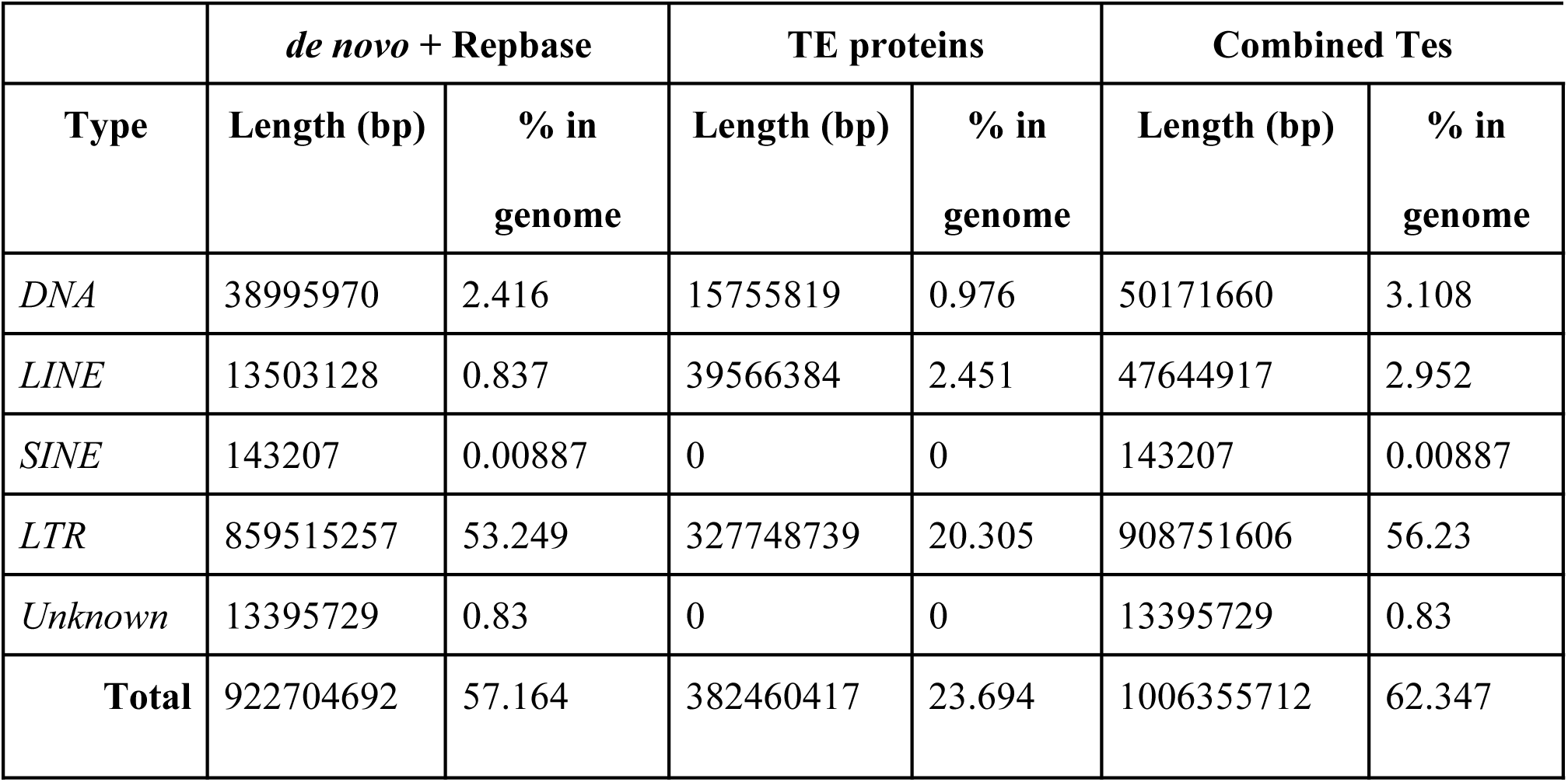
Detailed classification of repeat sequences identified in *Trochodendron aralioides. De novo* + Repbase: annotations predicted *de novo* by RepeatModeler, RepeatScout and LTR_FINDER; TE proteins: transposon elements annotated by RepeatMasker; Combined TEs: merged results from approaches above, with overlap removed. Unknown: repeat sequences RepeatMasker cannot classified.

The tRNA genes were identified by tRNAscan-SE [33] with the eukaryote set of parameters. The rRNA fragments were predicted by aligning them to *Arabidopsis thaliana* and *Oryza sativa* references rRNA sequences using BlastN (E-value of 1E-10). The miRNA and snRNA genes were predicted using INFERNAL [34] by searching against the Rfam database (release 9.1) (Supplementary Table S6).

### Genes annotation

#### RNA preparation and sequencing

RNA-seq was conducted for four tissue libraries (leaf, stem, bark, and bud) from the same individual as for the genome sequencing and assembly. A total of eight libraries were constructed (Supplementary Table S7). Total RNA was extracted using the RNAprep Pure Plant Kit (TIANGEN, Beijing, PR China) and gDNA contamination was removed with the RNase-Free DNase I (TIANGEN, Beijing, PR China). RNA quality was determined based on the estimation of the ratio of absorbance at 260nm/280nm (OD = 2.0) and the RIN (value = 9.2) by using a Nanodrop ND-1000 spectrophotometer (LabTech, USA) and a 2100 Bioanalyzer (Agilent Technologies, USA), respectively. The cDNA libraries were constructed with the NEBNext Ultra RNA Library Prep Kit for Illumina (New England Biolabs, Ipswich, Massachusetts, MA), following the manufacturer’s recommendations. Libraries were sequenced on an Illumina HiseqXTen platform (Illumina Inc., San Diego, CA), generating 150-bp paired-end reads. 26.7 Gb of clean RNA-seq sequences were produced, with at least 93.69% of the base with quality >Q20 (Supplementary Table S7).

RNA clean reads were both assembled into 312246 sequences using Trinity [35] and then annotated, and mapped against the genomic sequence using tophat v2.0.8 [36], then cufflinks v2.1.1 was used to assemble transcripts into gene models. Finally, all annotation results were combined with EVidenceModeler (EVM [38]) to obtain the final non-redundant gene set.

#### Annotation

Gene annotation was performed using three approaches:

- A homology-based approach, in which the protein sequences from *O. sativa, A. coerulea, F. excelsior, N. nucifera, Q. robur* and *V. vinifera* were aligned to the genome by using TblastN with an E-value cutoff by 1E-5. Blast hits were conjoined with software (BGI, Beijing, PR China). For each blast hit, Genewise [39] was used to predict the exact gene structure in the corresponding genomic regions.
- An *ab initio* gene prediction approach, using Augustus v2.5.5 [40], Genescan v1.0, GlimmerHMM v3.0.1 [41], Geneid [42] and SNAP [43] to predict coding genes on the repeats-masked *T. aralioides* genome.
- A transcriptome-based approach, in which RNA-seq data were mapped to genome (see above).

All gene models predicted from the above three approaches were combined by EVM into a non-redundant set of gene structures. Then we filtered out low quality gene models: (1) coding region lengths of ≤150 bp, (2) supported only by ab initial methods and with FPKM<1.

We identified an average of 5.1 exons per gene (mean length of 10.622 KB) in the *T. aralioides* genome (Table 1). The gene number, gene length distribution, CDS length distribution, exon length distribution and intron length distribution were all comparable to those of selected angiosperms species (Supplementary table S8, Supplementary Figure S3).

**Table 1.**
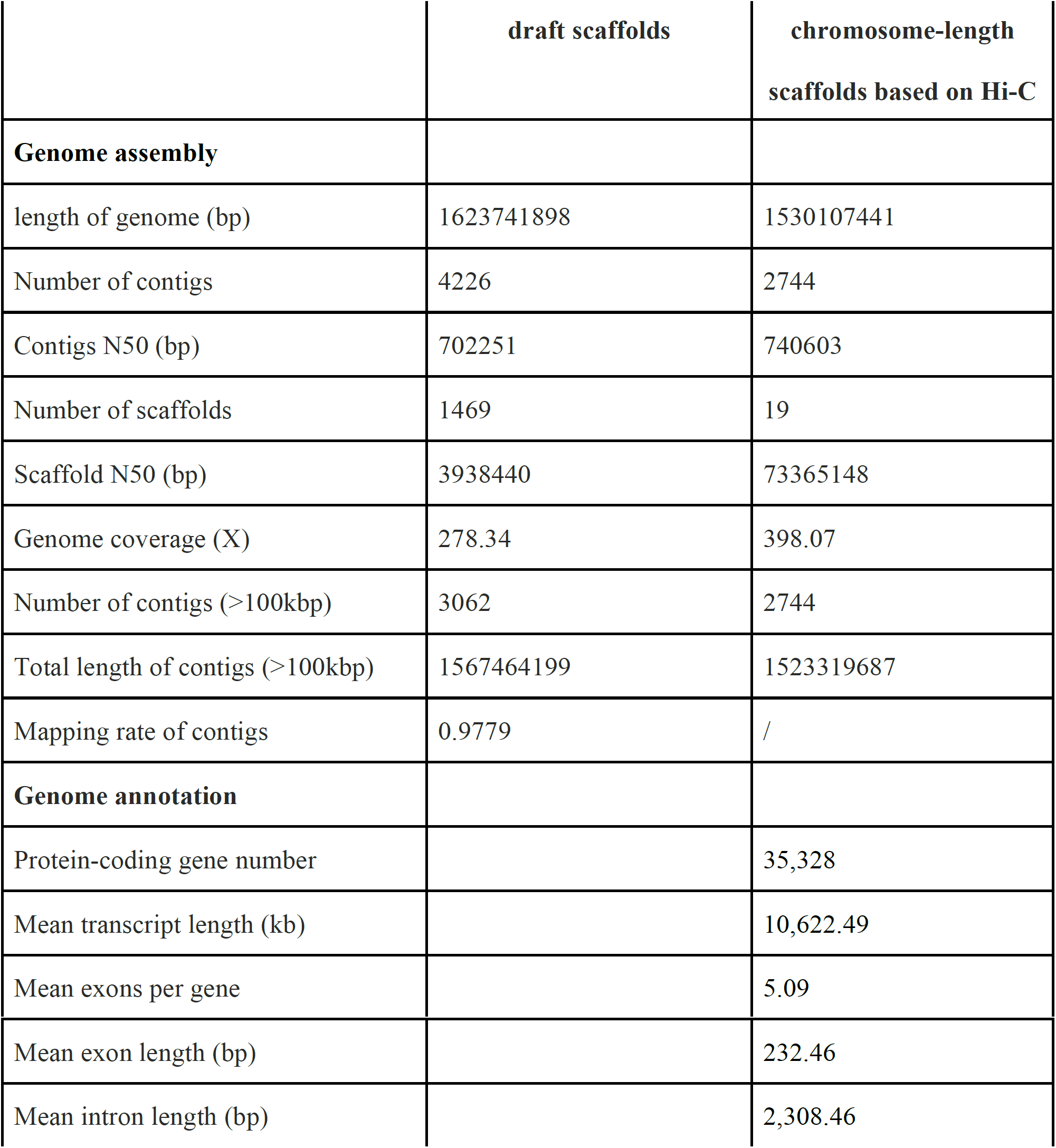
Summary of *Trochodendron aralioides* genome assembly and annotation.

Functional annotation was performed by blasting the protein coding genes sequences against SwissProt and TrEMBL using BLASTP (evalue 1E-05). The annotation information of the best BLAST hit from the databases, were transferred to our gene set annotations. Protein domains were annotated by searching InterPro(v32.0) and Pfam (V27.0) databases using InterProScan v4.8 [44] and Hmmer v3.1 [45], respectively. Gene Ontology (GO) terms for each gene were obtained from the corresponding InterPro or Pfam entry. The pathways in which the gene might be involved were assigned by blasting against the KEGG database (release53), with an E-value cutoff of 1E-05. The genes which were successfully annotated in GO, were classified into Biological process (BP), Cellular component (CC), Molecular Function (MF). Ultimately, 95.4% (33,696 genes) of the 35,328 genes were annotated by at least one database (Supplementary Table S9).

### Gene family identification and phylogenetic analyses of wheel-tree

Orthologous relationships between genes of *Amaranthus hypochondriacus, Amborella trichopoda, Annona muricata, Aquilegia coerulea, Arabidopsis thaliana, Helianthus annuus, Cinnamomum kanehirae, Musa acuminata, Nelumbo nucifera, Oryza sativa, Quercus robur* and *Vitis vinifera* were inferred through all-against-all protein sequence similarity searches with OrthoMCL [46] and only the longest predicted transcript per locus was retained(Supplementary Figure S4, S5).

For each gene family, an alignment was produced using Muscle [47], and ambiguously aligned positions trimmed using Gblocks [48]. A Maximum Likelihood (ML) tree was inferred using RAxML 7.2.9 [49].

Divergence times between species were calculated using MCMC, as implemented in PAML [50].

Nodes calibrations were defined from the TimeTree database (http://www.timetree.org/) as follows:

- the divergence between *Arabidopsis thaliana* and *Quercus robur* (97-109 Ma);
- the divergence between *Helianthus annuus* and *Amaranthus hypochondriacus* (107-116 Ma);
- the divergence between *Arabidopsis thaliana* and *Vitis vinifera* (109-114 Ma);
- the divergence between *Nelumbo nucifera* and *Vitis vinifera*(116-127 Ma);
- the divergence between *Musa acuminata* and *Oryza sativa* (90-115 Ma);
- the divergence between *Arabidopsis thaliana* and *Oryza sativa* (140-200 Ma);
- the divergence between *Amborella Trichopoda* and *Oryza sativa* (168-194 Ma).

The basal eudicot grade’s most basal representative, namely *A. coerulea*, diverged from other angiosperms during the lower Cretaceous 130.2 Ma (126.6-136.2 Ma), while *T. aralioides*, the most recently diverged basal eudicot, diverged from the core-eudicots approximately 124 Ma (121.8-125.8 Ma). Finally, the divergence between rosids and asterids, and thus the crown age of core eudicots was reconstructed as 114.0 Ma (111.3-116.2 Ma).

To identify gene families that experienced a significant expansion or contraction during the evolution of the wheel-tree, we used the likelihood model implemented in CAFE [51], with default parameters. The phylogenetic tree topology and branch lengths were taken into account to infer the significance of change in gene family size in each branch (Figure 4). The genes families that experienced the most significant expansions were mainly involved in pathogen/stress response [e.g. the cyanoamino-acid metabolism, p=2.35×10^−28^; the plant-pathogen interaction map, p=2.29×10^−22^; the tryptophan metabolism, p=5.85×10^−10^] (Supplementary Table S10).

**Figure 3.**
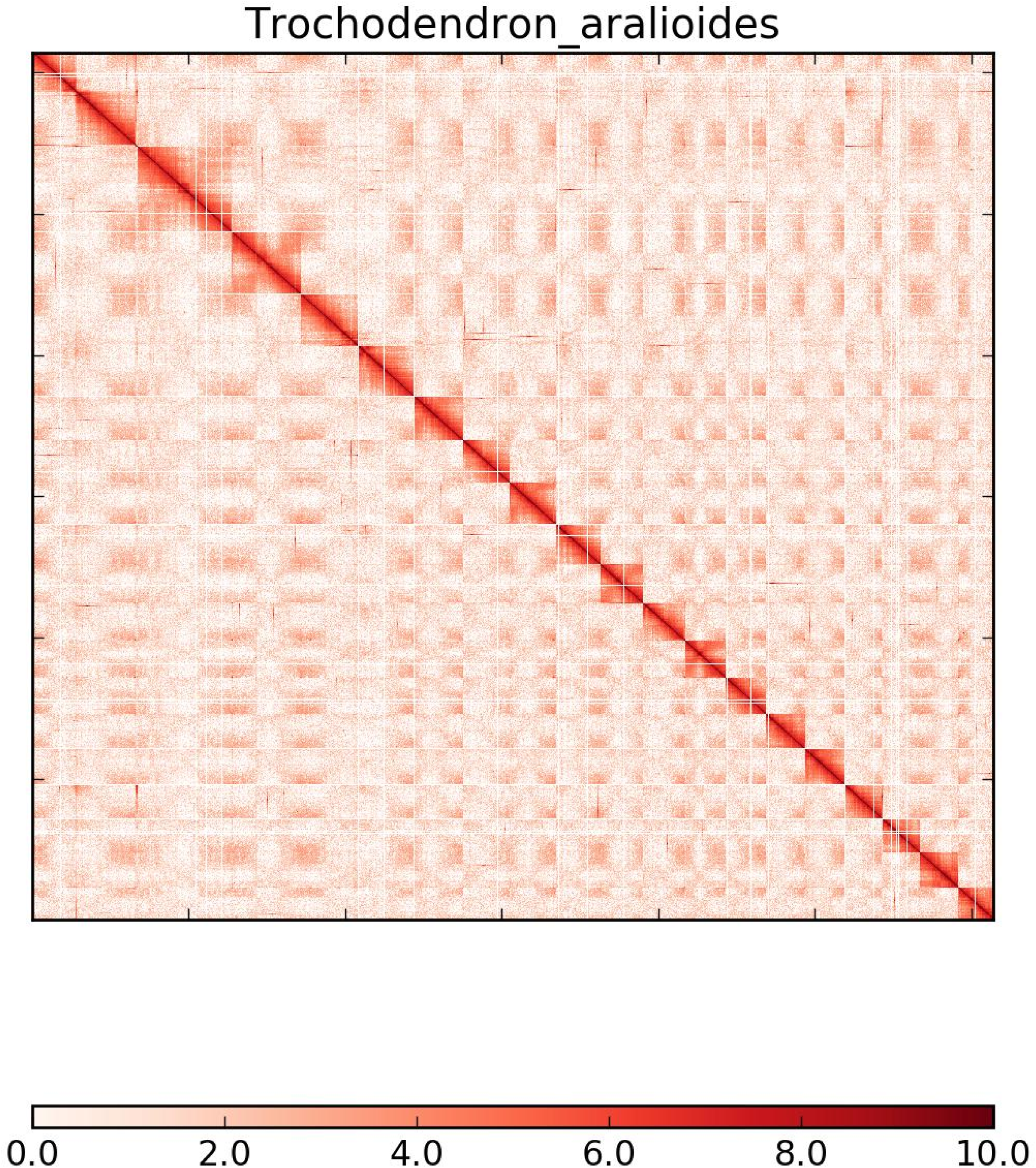
Hi-C interaction heat map for *T. aralioides* reference genome showing interactions between the 19 chromosomes.

**Figure 4.**
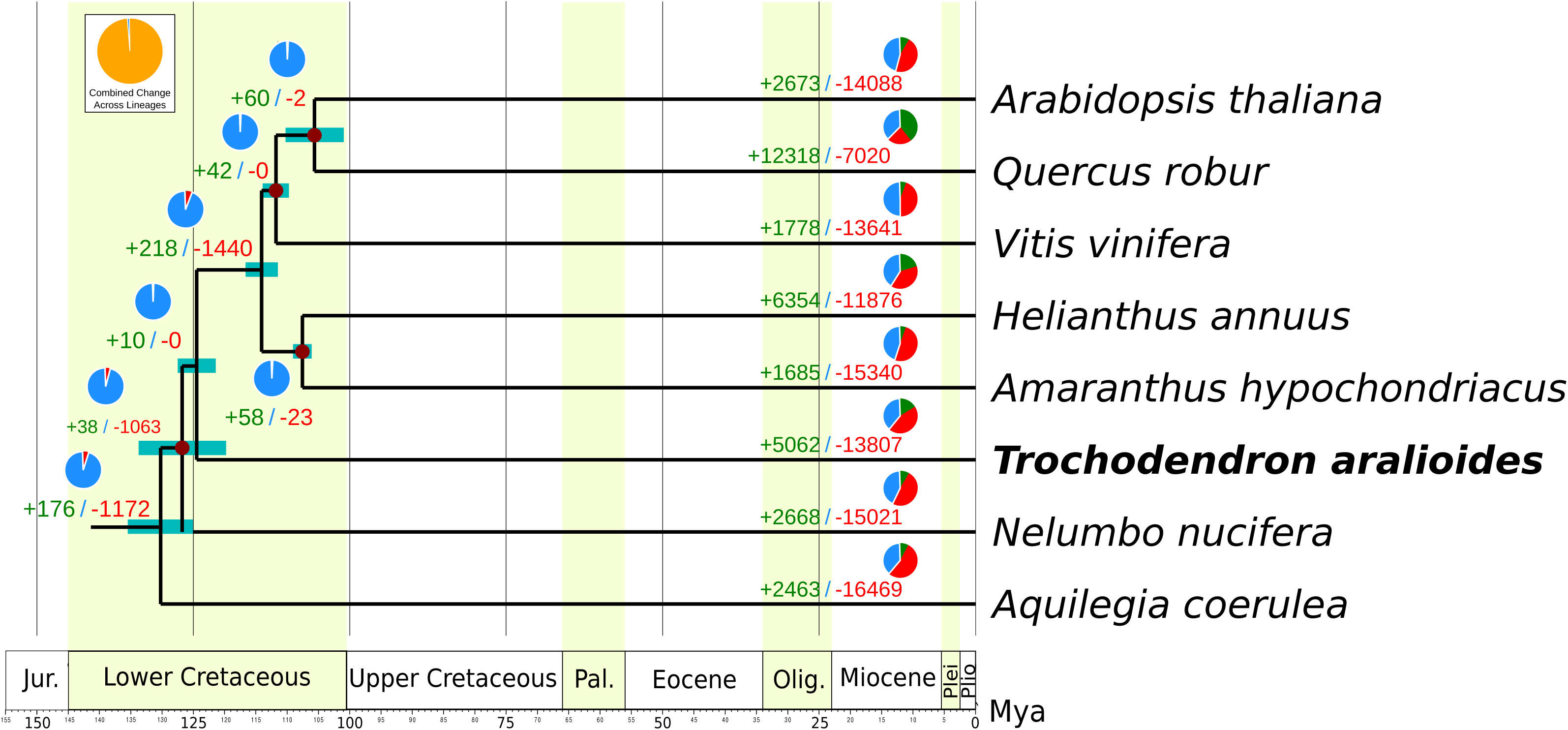
Phylogenetic tree and number of gene families displaying expansion and contraction among 12 plant species. The pie charts show the expansion (green), contraction (red) and conserved (blue) gene family proportions among all gene families. Estimated divergence time confidence interval are shown at each internal node as teal bars. Calibrated nodes indicated by red dots (see text for details on calibration scheme).

### Positive selection

Positive selection is a major driver of biological adaptation. Using the protein-coding sequences, we calculated the number of synonymous substitutions per site (Ks) and nonsynonymous substitutions per site (Ka) and assessed the deviation from zero of the difference Ka − Ks. A Ka/Ks > 1 represent an evidence of positive selection. We calculated the nonsynonymous to synonymous substitution rate ratio (dN/dS), which is also commonly interpreted as evidence for positive selection. MUSCLE [47] was used to align the protein and nucleotide sequences, then we used Gblocks [48] to eliminate poorly aligned positions and divergent regions from the alignment. The maximum likelihood-based branch lenghths test of PAML package [50] was used for the comparisons, and the ratio Ka/Ks was calculated over the entire length of the protein coding gene. Using *T. aralioides* as the foreground branch and *Amaranthus hypochondriacus, Helianthus annuus, Nelumbo nucifera, Aquilegia coerulea* as background branches, we identified 238 genes were considered as candidate genes under positive selection (p-value<0.01, FDR < 0.05) using a maximum likelihood-based branch lengths test. A GO and KEGG enrichment analysis showed that positive selection was especially detected in cell metabolism (such as vitamins and amino-acid biosynthesis, Supplementary Table S11).

### Whole-genome duplication analysis

Mcscan [52] was used to identify collinear segments within *Trochodendron* and between the *T. aralioides* and other angiosperm genomes. The sequences of the gene pairs contained in the (inter-) collinear segments of the genome were extracted and the *codeml* tool in the PAML package [50] was used to calculate the value of 4dTv. The distribution of 4dTv can reflect whether genome-wide replication events occur in the evolutionary history of species, the relative time of genome-wide replication events, and the divergent events among species.

The 4dTv values of all paralogous gene pairs in *Trochodendron aralioides* were calculated, as well as those in *Aquilegia coerulea, Helianthus annuus* and *Annona muricata* for comparisons. In addition, the 4dTv values were calculated for all ortholog gene pairs between *Trochodendron aralioides* and *A. coerulea, H. annuus* and *A. muricata* to observe species divergence events. The peak of 4dTv distribution in the *T. aralioides* genome was around ∼0.1, whereas the peak for interspecific comparisons with *A. coerulea* and *H. annuus* were around ∼0.3 and ∼0.2, respectively (Supplementary Figure S6). All together, these 4dTv distributions indicate that a whole genome replication event occurred in *T. aralioides* after its divergence from both other basal angiosperms and core-eudicots.

## Conclusions

We successfully assembled the genome of *T. aralioides* and report the first chromosome-level genome sequencing, assembly and annotation based on long reads from the third-generation PacBio Sequel sequencing platform for basal eudicotyledons. The final draft genome assembly is approximately 1.614 Gb, which is slightly smaller than both the estimated genome size (1.758 Gb) based on k-mer analysis and on cytometry (1.868 Gb, [16]). With a contig N50 of 691 Kb and a scaffold N50 of 73.37 Mb, the chromosome-level genome assembly of *T. aralioides* is the first high-quality genome in the Trochodendrales order. We also predicted 35,328 protein-coding genes from the generated assembly, and 95.4% (33,696 genes) of all protein-coding genes were annotated. We found that the divergence time between *T. aralioides* and its common ancestor with the core eudicots was approximately 124.2 Ma. The chromosome-level genome assembly together with gene annotation data generated in this work will provide a valuable resource for further research on floral morphology diversity, on the early evolution of eudicotyledons and on the conservation of this iconic tree species.

## Availability of supporting data

Supporting data and materials are available in the GigaScience GigaDB database [53] (***pending***), and raw sequences deposited in the EBI database under the accession number (***pending***).

## Abbreviation

BUSCO: Benchmarking Universal Single-Copy Orthologs;
CDS: Coding sequence;
GO: Gene ontology;
KEGG: Kyoto Encyclopaedia of Genes and Genomes;
LINE: Long interspersed nuclear elements;
LTR: Long terminal repeats;
NGS: Next Generation Sequencing;
TE: Transposable elements.

## Supplementary Materials

**Supplementary Figure S1.** GC content analysis of *T. aralioides* genome based on Illumina reads for genome size survey.

**Supplementary Figure S2.** Divergence distribution of transposable elements in the genome of *T. aralioides*. Units in Kimura substitution level (CpG adjusted).

**Supplementary Figure S3.** Genes characteristics in *T. aralioides* and other angiosperms. From left to right and top to bottom: lengths of messenger RNA; lengths of exons in coding regions; number of exons per gene; lengths of introns in genes; lengths of coding regions (CDS). Aco: *Aquilegia caeruleus*; Fex: *Fraxinus excelsior*; Nun: *Nelumbo nucifera*; Osa: *Oryza sativa*; Qro: *Quercus robur*; Tar: *Trochondendron aralioides*; Vvi: *Vitis vinifera*.

**Supplementary Figure S4.** Comparing orthogroups between *T. aralioides* and other angiosperms species. Aco: *Aquilegia caeruleus*; Ahy: *Amaranthus hypochondriacus*; Amu: *Annona muricata;* Ath: *Arabidopsis thaliana*; Atr: *Amborella Trichopoda*; Cka: *Cinnamomum micranthum*; Han: *Helianthus annuus*; Mac: *Musa acuminata*; Nnu: *Nelumbo nucifera*; Osa: *Oryza sativa*; Qro: *Quercus robur*; Tar: *Trochondendron aralioides*; Vvi: *Vitis vinifera*.

**Supplementary Figure S5.** Orthologous gene families across four angiosperms genomes (*Trochodendron aralioides, Annona muricata, Amaranthus hypochondriacus* and *Aquilegia coerulea*).

**Supplementary Figure S6.** Orthologous gene families across four angiosperms genomes (*Trochodendron aralioides, Annona muricata, Amaranthus hypochondriacus* and *Aquilegia coerulea*).

