## Supplementary figures and images for "*Trochodendron aralioides*, the first chromosome-level draft genome in Trochodendrales and a valuable resource for basal eudicot research"

### Supplementary Figure 1

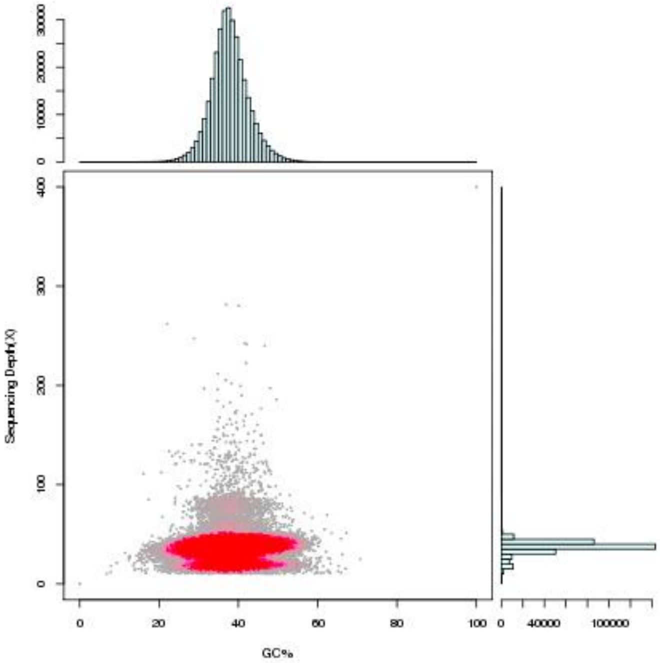

### Supplementary Figure 2

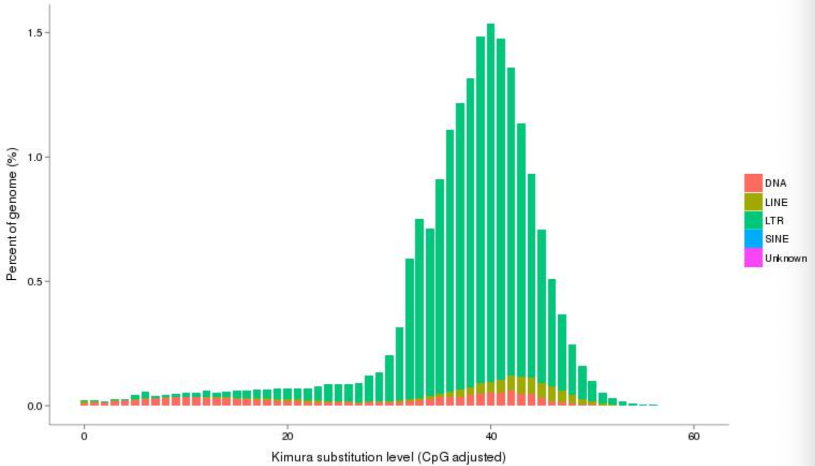

### Supplementary Figure 3

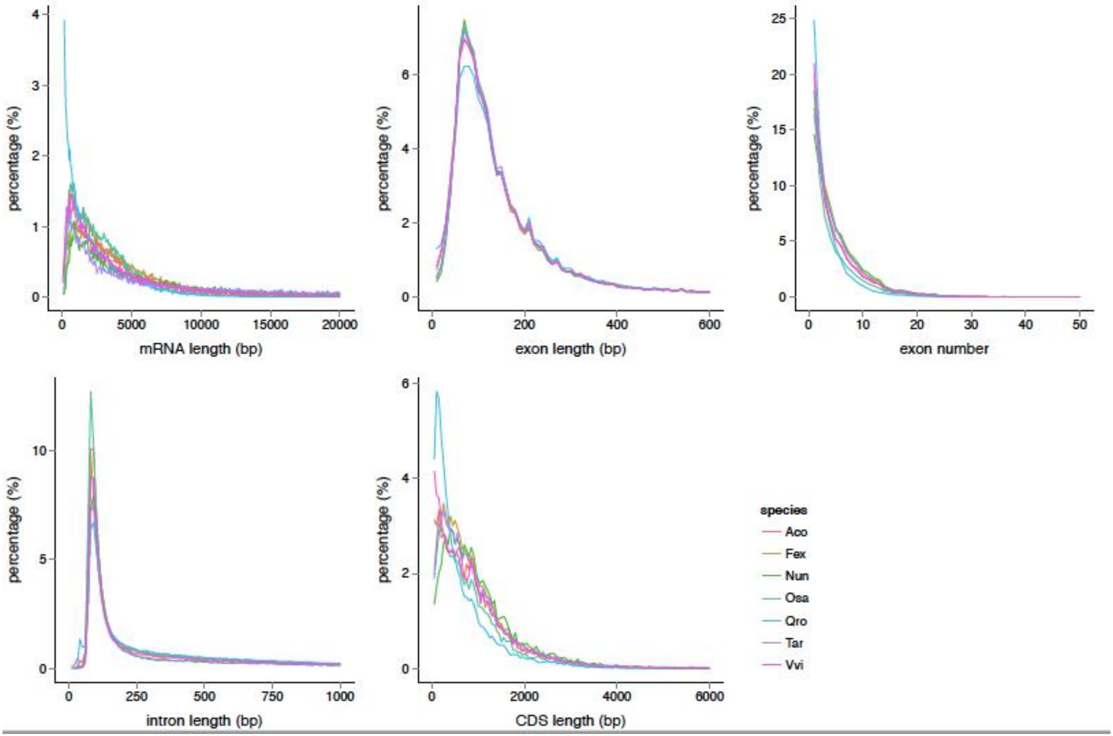

### Supplementary Figure 4

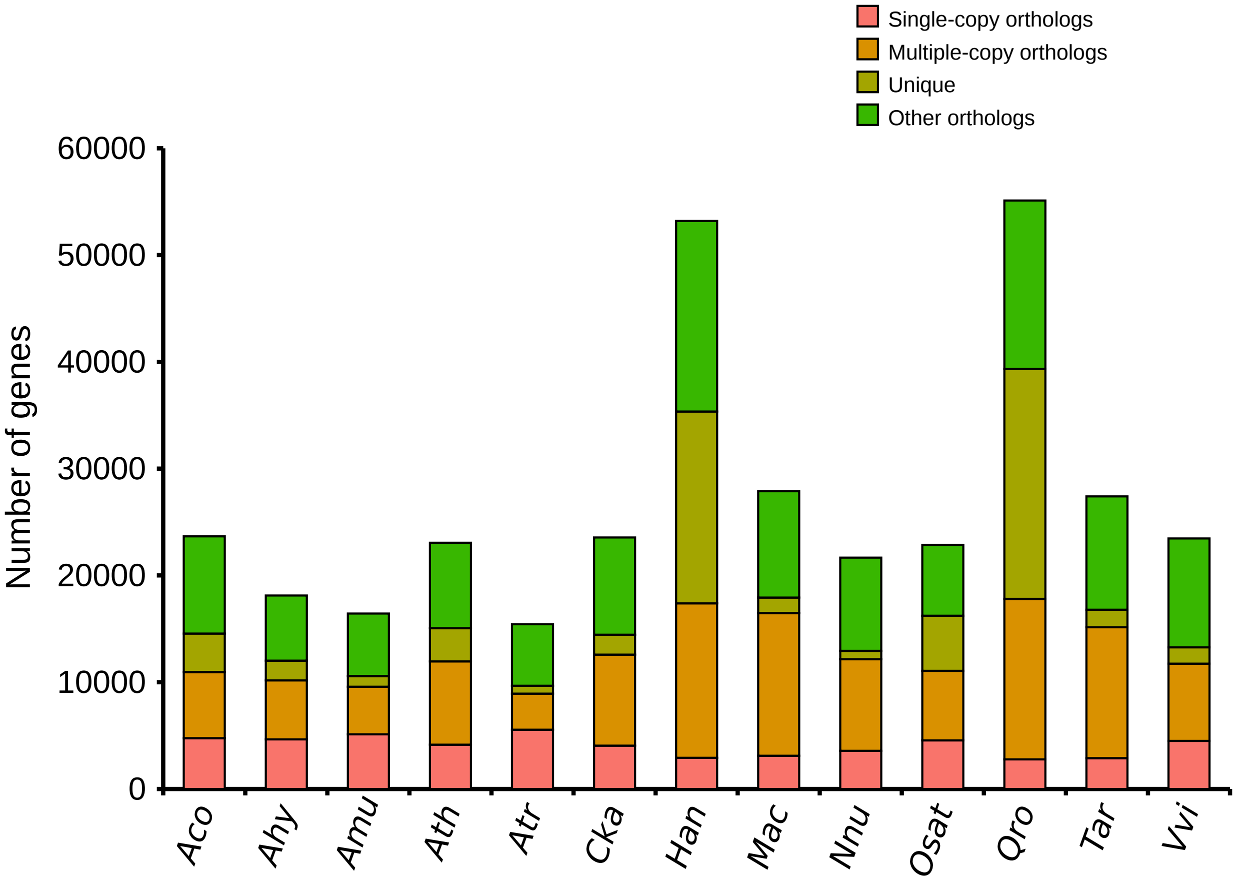

### Supplementary Figure 5

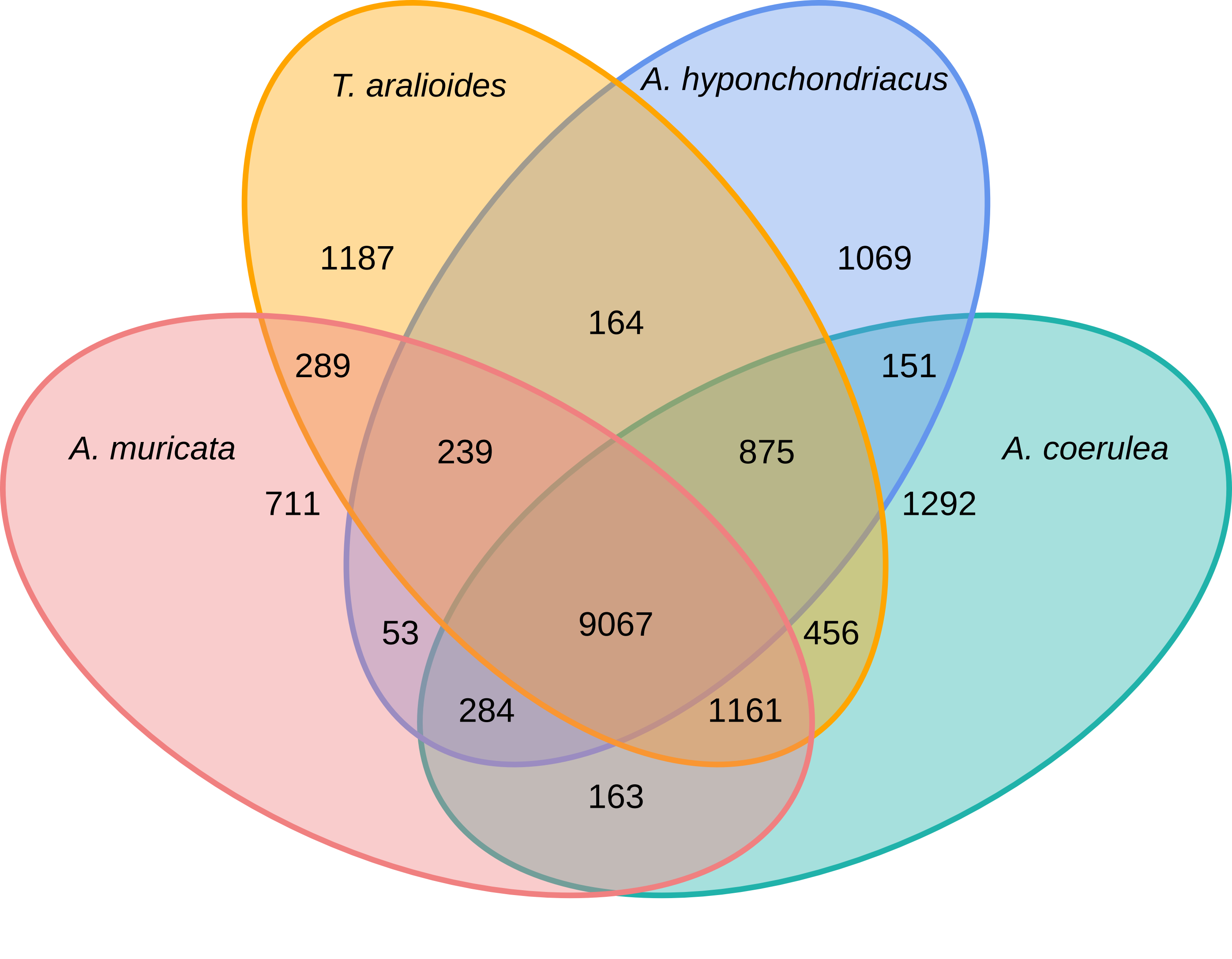

### Supplementary Figure 6

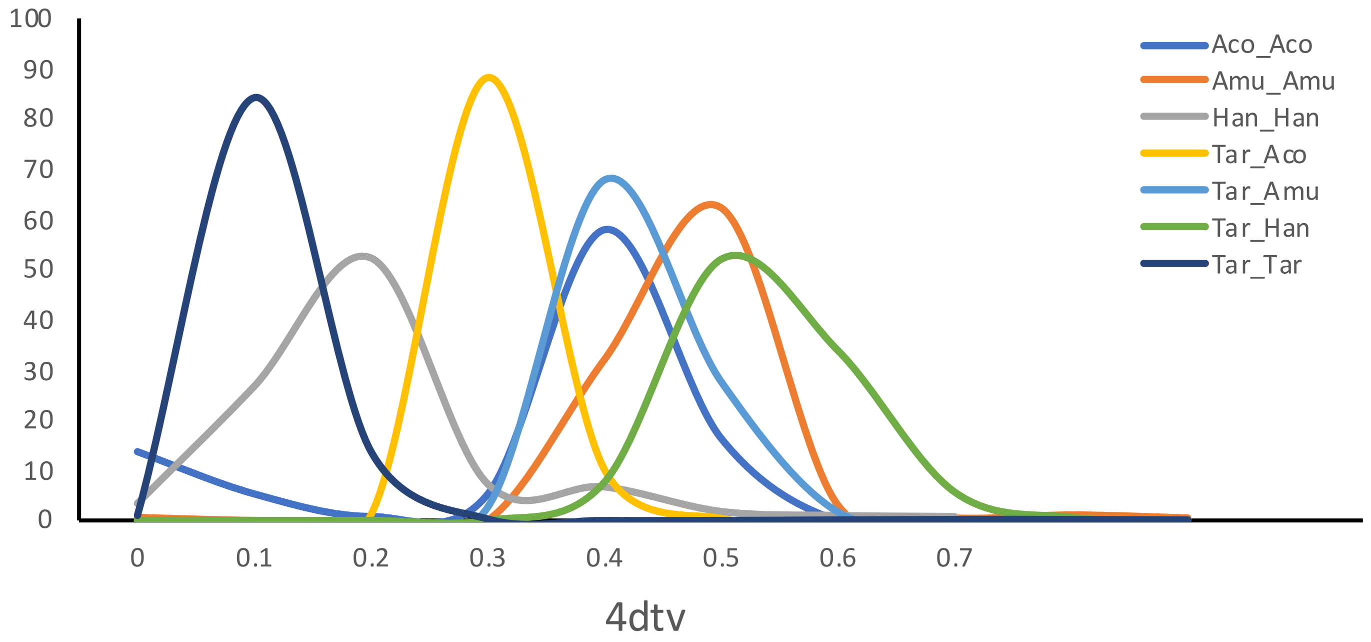
