## Supplementary Tables for "*Trochodendron aralioides*, the first chromosome-level draft genome in Trochodendrales and a valuable resource for basal eudicot research"

**Supplementary Table S1** Summary of sequence data from *T. aralioides.*

| Type | Method | Library size (bp) | Data size (clean Gb) | Read N50 (bp) |
|---|---|---|---|---|
| DNA | NovaSeq | 300~350 | 124.33 | 150 |
| DNA | PacBio Sequel | 20 000 | 177.80 | 13,171 |
| DNA (10X Genomics) | NovaSeq | 300~350 | 183.52 | 150 |
| DNA (Hi-C) | NovaSeq | 300~350 | 193.90 | 150 |
| RNA | NovaSeq | 150 | 26.7 | 150 |
| | | **Total:** | 679.55 | |

**Supplementary Table S2** Sequencing quality assessment of Hi-C sequencing data.

| Sample | Raw Base(bp) | Clean Base(bp) | Effective **Rate (%)** | Error Rate(%) | Q20(%) | Q30(%) | GC content(%) |
|---|---|---|---|---|---|---|---|
| RHC00944-1_L4 | 17,736,454,200 | 16,815,348,300 | 94.81 | 0.02 | 97.42 | 94.31 | 41.52 |
| RHC00944-1_L5 | 25,606,849,200 | 24,351,238,500 | 95.1 | 0.02 | 97.2 | 93.83 | 41.48 |
| RHC00944-1_L6 | 37,548,328,800 | 35,089,588,500 | 93.45 | 0.03 | 96.66 | 92.22 | 41.53 |
| RHC00944-1_L8 | 129,596,693,700 | 117,647,530,500 | 90.78 | 0.03 | 95.2 | 89.45 | 41.61 |

**Supplementary Table S3** Genome quality of *T. aralioides* based on the BUSCO and CEGMA assessments.

| | BUSCO | | CEGMA | |
|---|---|---|---|---|
| **Type** | **Proteins** | **Percentage (%)** | **Proteins** | **Percentage (%)** |
| Complete (BUSCO and CEGMA) | 1,316 | 91.4 | 216 | 87.10 |
| Complete and single-copy (BUSCO) | 1,096 | 76.1 | | |
| Complete and duplicated (BUSCO) | 220 | 15.3 | | |
| Fragmented (BUSCO) | 40 | 2.8 | | |
| Complete and partial (CEGMA) | | | 232 | 93.55 |
| Missing (BUSCO) | 84 | 5.8 | | |
| Total groups searched (BUSCO and CEGMA) | 1,440 | | 248 | |

**Supplementary Table S4** Chromosome length distribution in *T. aralioides*, as inferred by Lachesis using Hi-C sequencing data.

| Chr_id | Length |
|---|---|
| Lachesis_group0__255_contigs__length_166831109 | 167,510,055 |
| Lachesis_group1__235_contigs__length_150294801 | 150,813,425 |
| Lachesis_group2__174_contigs__length_109165540 | 109,591,199 |
| Lachesis_group3__190_contigs__length_91497359 | 91,953,593 |
| Lachesis_group4__147_contigs__length_87728453 | 88,019,702 |
| Lachesis_group5__98_contigs__length_77627035 | 77,886,091 |
| Lachesis_group6__184_contigs__length_74161915 | 74,576,108 |
| Lachesis_group7__131_contigs__length_73030964 | 73,365,148 |
| Lachesis_group8__137_contigs__length_70371694 | 70,682,679 |
| Lachesis_group9__129_contigs__length_67714956 | 67,974,108 |
| Lachesis_group10__152_contigs__length_65993342 | 66,344,902 |
| Lachesis_group11__124_contigs__length_64544952 | 64,834,175 |
| Lachesis_group12__120_contigs__length_64144231 | 64,446,891 |
| Lachesis_group13__129_contigs__length_61810036 | 62,138,229 |
| Lachesis_group14__65_contigs__length_62905247 | 63,102,812 |
| Lachesis_group15__130_contigs__length_59618705 | 59,906,625 |
| Lachesis_group16__142_contigs__length_58914604 | 59,218,580 |
| Lachesis_group17__77_contigs__length_60868391 | 61,055,640 |
| Lachesis_group18__125_contigs__length_56368853 | 56,687,479 |
| **TOTAL** | **1,530,107,441 (94.8%)** |

**Supplementary Table S5** Final *T. aralioides* assembly information.

| | **Input Assembly** | **LACHESIS assembly** |
|---|---|---|
| Total Length | 1,623.74Mb | 1,614.13Mb |
| L50/N50 | 92 Scaffolds; 3.94Mb | 8 Scaffolds; 73.37Mb |
| L90/N90 | 458 Scaffolds; 785.85 kb | 18 Scaffolds; 59.22 Mb |
| Longest Scaffolds | 57.56 Mb | 167.51 Mb |
| Number of Scaffolds | 1,469 | 1,534 |

| | | |
|---|---|---|
| Contig N50 | 702.25 Kb | 691.19 Kb |

**Supplementary Table S6** Statistics of the annotation of non-coding RNAs in the *T. aralioides* genome.

| Type | | Copy | Average length (bp) | Total length (bp) | % of genome |
|---|---|---|---|---|---|
| **miRNA** | | 1,536 | 132.52 | 203,547 | 0.013 |
| **tRNA** | | 870 | 74.79 | 64,986 | 0.004 |
| **rRNA** | **rRNA** | 198 | 228.99 | 45,340 | 0.0028 |
| | **18S** | 71 | 440.25 | 31,258 | 0.0019 |
| | **28S** | 51 | 112.49 | 5,737 | 0.00036 |
| | **5.8S** | 19 | 103.15 | 1,960 | 0.00012 |
| | **5S** | 57 | 112.01 | 6,385 | 0.000396 |
| **snRNA** | **snRNA** | 1,151 | 111.96 | 128,864 | 0.00798 |
| | **CD-box** | 810 | 104.72 | 84,827 | 0.00526 |
| | **HACA-box** | 90 | 127.06 | 11,435 | 0.00071 |
| | **Splicing** | 250 | 129.57 | 32,392 | 0.002 |

**Supplementary Table S7** Library construction details.

| Sample | Raw Reads | Clean Reads | Clean Bases | Error(%) | Q20(%) | Q30(%) | GC Content(%) | Nb Gene | Read N50 | Max length | Average length |
|---|---|---|---|---|---|---|---|---|---|---|---|
| leaves 1 | 23424919 | 21967266 | 3.3G | 0.02 | 98.31 | 95.37 | 45.86 | 234694 | 1235 | 13804 | 696 |
| leaves 2 | 23424919 | 21967266 | 3.3G | 0.04 | 94.15 | 86.68 | 45.66 | | | | |
| bark 1 | 22622608 | 21144982 | 3.17G | 0.02 | 98.26 | 95.26 | 46.27 | | | | |
| bark 2 | 22622608 | 21144982 | 3.17G | 0.04 | 93.9 | 86.18 | 46.09 | | | | |
| buds 1 | 24260878 | 22773120 | 3.42G | 0.02 | 98.28 | 95.32 | 45.73 | | | | |

| | | | | | | | |
|---|---|---|---|---|---|---|---|
| buds 2 | 24260878 | 22773120 | 3.42G | 0.04 | 93.69 | 85.81 | 45.52 |
| stems 1 | 24467354 | 22927722 | 3.44G | 0.02 | 98.29 | 95.33 | 45.84 |
| stems 2 | 24467354 | 22927722 | 3.44G | 0.04 | 93.8 | 86 | 45.64 |

**Supplementary Table S8** Gene annotation of the *T. aralioides* genome.

| Gene set | | Number | Average gene length (bp) | Average CDS length (bp) | Average exon per gene | Average exon length (bp) | Average intron length (bp) |
|---|---|---|---|---|---|---|---|
| ***ab initio*** | *AUGUSTUS* | 51,969 | 7,124.24 | 971.33 | 3.81 | 254.9 | 2,189.12 |
| | *Glimmer HMM* | 123,116 | 11,693.65 | 488.14 | 2.85 | 171.36 | 6,061.3 |
| | *SNAP* | 36,880 | 17,703.12 | 495.31 | 3.3 | 150.24 | 7,492.35 |
| | *Genscan* | 77,247 | 13,712.45 | 949.08 | 5.55 | 170.95 | 2,803.95 |
| | *Geneid* | 128,396 | 4,670.57 | 573.15 | 3.53 | 162.58 | 1,622.57 |
| **Homology** | *Oryza sativa* | 64,868 | 3,277.66 | 1,358.74 | 2.36 | 575.19 | 1,408.63 |
| | *Aquilegia coerulea* | 44,155 | 2,797 | 655.21 | 2.21 | 297.07 | 1,776.61 |
| | *Fraxinus excelsior* | 48,727 | 3,416.48 | 764.78 | 2.42 | 315.84 | 1,865.54 |
| | *Nelumbo nucifera* | 32,584 | 6,832.52 | 1,262.44 | 3.67 | 344.41 | 2,089.64 |
| | *Quercus robur* | 109,500 | 2,519.72 | 909.75 | 2.1 | 433.64 | 1,466.4 |
| | *Vitis vinifera* | 64,058 | 4,148.63 | 926.06 | 3.06 | 302.28 | 1,561.64 |
| **RNAseq** | *Cufflinks* | 61,576 | 17,349.81 | 2,060.41 | 6.13 | 335.91 | 2,978.17 |
| | *PASA* | 44,888 | 9,093.64 | 972.02 | 4.38 | 222.1 | 2,405.28 |
| EVM | | 53,020 | 8,187.19 | 972.59 | 4.13 | 235.35 | 2,303.16 |
| PASA-update * | | 52,819 | 8,235.28 | 975.9 | 4.13 | 236.52 | 2,322.22 |
| Final set * | | 35,328 | 10,622.49 | 1,183.03 | 5.09 | 232.46 | 2,308.46 |

* contain UTR region.

**Supplementary Table S9** Functional annotation of the protein-coding genes in *T. aralioides* genome.

| Type | Number (%) | Percent (%) |
|---|---|---|
| Total | 35,328 | |

| | | |
|---|---|---|
| InterPro | 29,122 | 82.4 |
| GO | 19,982 | 56. |
| KEGG | 26,318 | 74.5 |
| Pfam | 27,101 | 76.7 |
| Swissprot | 27,659 | 78.3 |
| NR | 33,625 | 95.2 |
| Annotated | 33,696 | 95.4 |
| Unannotated | 1,632 | 4.6 |

**Supplementary Table S10** The 20 top KEGG pathways annotations for genes families that experienced copy number expansions.

| MapID | MapTitle | AdjustedPv |
|---|---|---|
| map00460 | Cyanoamino acid metabolism | 2.35168663154486E-28 |
| map05164 | Influenza A | 2.35168663154486E-28 |
| map04113 | Meiosis - yeast | 3.47267639636242E-24 |
| map04626 | Plant-pathogen interaction | 2.28605340320955E-22 |
| map00908 | Zeatin biosynthesis | 1.78710186484916E-16 |
| map05162 | Measles | 1.13150653970762E-14 |
| map00830 | Retinol metabolism | 8.14983642460509E-14 |
| map00380 | Tryptophan metabolism | 5.84909900425868E-10 |
| map00940 | Phenylpropanoid biosynthesis | 5.84909900425868E-10 |
| map00350 | Tyrosine metabolism | 7.97070204554834E-10 |
| map00970 | Aminoacyl-tRNA biosynthesis | 0.000000001 |
| map00402 | Benzoxazinoid biosynthesis | 1.13959596153542E-09 |
| map00950 | Isoquinoline alkaloid biosynthesis | 3.19244057770548E-09 |
| map00941 | Flavonoid biosynthesis | 1.84819219426448E-08 |
| map04728 | Dopaminergic synapse | 2.86756991206953E-08 |
| map04722 | Neurotrophin signaling pathway | 3.25903616565627E-08 |
| map04064 | NF-kappa B signaling pathway | 6.93633061727493E-07 |
| map05140 | Leishmaniasis | 7.55346618276399E-07 |
| map05142 | Chagas disease (American trypanosomiasis) | 7.55346618276399E-07 |
| map05217 | Basal cell carcinoma | 7.55346618276399E-07 |

**Supplementary Table S11** The 20 KEGG pathways under positive selection.

| MapID | MapTitle | AdjustedPv |
|---|---|---|
| map04215 | Apoptosis - multiple species | 0.278326703 |
| map03013 | RNA transport | 0.5687306488 |
| map00740 | Riboflavin metabolism | 0.5687306488 |
| map04122 | Sulfur relay system | 0.5687306488 |
| map04146 | Peroxisome | 0.5687306488 |
| map04919 | Thyroid hormone signaling pathway | 0.5687306488 |
| map00240 | Pyrimidine metabolism | 0.5687306488 |
| map03440 | Homologous recombination | 0.5687306488 |
| map00563 | Glycosylphosphatidylinositol (GPI)-anchor biosynthesis | 0.6287520051 |
| map05152 | Tuberculosis | 0.6450823973 |
| map00590 | Arachidonic acid metabolism | 0.6450823973 |
| map00230 | Purine metabolism | 0.6450823973 |
| map03450 | Non-homologous end-joining | 0.6450823973 |
| map03022 | Basal transcription factors | 0.6450823973 |
| map05217 | Basal cell carcinoma | 0.6450823973 |
| map03020 | RNA polymerase | 0.6450823973 |
| map04626 | Plant-pathogen interaction | 0.6450823973 |
| map00400 | Phenylalanine, tyrosine and tryptophan biosynthesis | 0.6450823973 |
| map04977 | Vitamin digestion and absorption | 0.6737771431 |
| map00053 | Ascorbate and aldarate metabolism | 0.6737771431 |
